# CameraTrapDetectoR: Automatically detect, classify, and count animals in camera trap images using artificial intelligence

**DOI:** 10.1101/2022.02.07.479461

**Authors:** Michael A. Tabak, Daniel Falbel, Tess Hamzeh, Ryan K. Brook, John A. Goolsby, Lisa D. Zoromski, Raoul K. Boughton, Nathan P. Snow, Kurt C. VerCauteren, Ryan S. Miller

## Abstract

Motion-activated wildlife cameras, or camera traps, are widely used in biological monitoring of wildlife. Studies using camera traps amass large numbers of images and analyzing these images can be a large burden that inhibits research progress. We trained deep learning computer vision models using data for 168 species that automatically detect, count, and classify common North American domestic and wild species in camera trap images. We provide our trained models in an R package, CameraTrapDetectoR. Three types of models are available: a taxonomic class model classifies objects as mammal (human and non-human) or avian; a taxonomic family model that recognizes 31 mammal, avian, and reptile families; a species model that recognizes 75 domestic and wild species including all North American wild cat species, bear species, and Canid species. Each model also includes a category for vehicles and empty images. The models performed well on both validation datasets and out-of-distribution testing datasets as mean average precision values ranged from 0.80 to 0.96. CameraTrapDetectoR provides predictions as an R object (a data frame) and flat file and provides the option to create plots of the original camera trap image with the predicted bounding box and label. There is also the option to apply models using a Shiny Application, with a point-and-click graphical user interface. This R package has the potential to facilitate application of deep learning models by biologists using camera traps.

## Introduction

Camera traps, motion-activated field cameras, are often used in studies of wildlife or for routine monitoring and have facilitated great advancements in ecological research and wildlife monitoring (Jumeau et al. 2017, O’Connor et al. 2017). Camera trap data has become a foundational data source for estimating population size, changes in population, species diversity, animal movement, and inter- and intra-species competition (Sollmann 2018). However, camera traps can accumulate millions of images, that require evaluation to identify animals or extract other biological information (Swanson et al. 2015, Niedballa et al. 2016). The burden of manually reviewing camera trap images has limited the broad scale use for landscape scale monitoring and frequently limits the number of cameras and the length of time cameras are deployed in research studies. The burden of manually reviewing camera trap images has prompted efforts to automatically analyze images using deep learning computer vision methods (Willi et al. 2019, Schneider et al. 2020). Image classification is a deep learning computer vision technique that classifies the entire image as one or more categories (He et al. 2016), and has shown to be effective at classifying some datasets of camera trap images (Norouzzadeh et al. 2018, Tabak et al. 2019). However, when image classification models are deployed on out-of-distribution datasets (i.e., datasets from camera traps not included in the training data), they often perform poorly at recognizing animal species (Schneider et al. 2020, Tabak et al. 2020b). The poor performance of image classification models in out-of-distribution datasets has prompted the use of object detection models to analyze camera trap images.

Object detection models find and classify regions of interest in images (Zou et al. 2019). When applied to camera trap image datasets, object detection algorithms can locate animals in an image, identify each species of animal in an image, and count the number of animals by species (Norouzzadeh et al. 2021). Object detection has been shown to be more effective than whole image classification for finding empty images (those without animals) and accurately classifying animals (Beery et al. 2019, 2020). Currently, software options exist for applying object detection algorithms to camera trap images, specifically, Wildlife Insights (Wildlife Insights 2021), MegaDetector (Beery et al. 2019), Conservation AI (Chalmers et al. 2019), AnimalFinder (Price Tack et al. 2016), and ClassifyMe (Falzon et al. 2020). Each of these methods have strengths and limitations, which are discussed in detail in a recent review of these software packages (Vélez et al. 2022). Despite the availability of detection algorithms for identifying animals in camera trap images an R based tool is currently not available that can be integrated into workflows.

Our goal is to provide an object detection option that draws from the strengths of other software packages while improving functionality for ecologists and on-the-ground wildlife management, making automatic wildlife imagery processing easily obtainable. Specifically, we take a taxonomic approach allowing identification and classification of images into taxonomic groups: class, family, and species. This can be particularly useful if the species of interest is not included in the species level model. Additionally, we sought to develop a tool that can be easily incorporated into automated or semi-automated workflows, particularly for ecological monitoring applications. Lastly, because camera trap data frequently has data privacy concerns our R package has increased utility because can be deployed locally on a laptop computer and does not require a graphics processing unit (GPU). We also provide a Shiny Application (Chang et al. 2021) that allows users to deploy models with a point-and-click graphical user interface. The R package introduced here, CameraTrapDetectoR, is freely available from https://github.com/TabakM/CameraTrapDetectoR.

## Methods

### Camera trap images and annotations

We used select images from the North American Camera Trap Images (NACTI) dataset (Tabak et al. 2019). Images were selected from the NACTI dataset using a conditional Latin hypercube sampling approach accounting for study site and camera number. This ensured that images used for training and validation were evenly distributed across both studies and camera sites to account for potential differences. While the NACTI database does have labels for species and vehicles it does not contain bounding boxes identifying the location of animals in each image. For images containing animals or vehicles, we drew bounding boxes around these objects using the MegaDetector (Beery et al. 2019). Each image was then manually reviewed to ensure the bounding box was accurate. We excluded images with bounding boxes that overlapped to limit any potential confounding with the object detection approach. The NACTI dataset does not contain images for some important North American wildlife species such as ocelot. To increase the usability of CameraTrapDetectoR we also manually generated bounding boxes using images obtained from Labeled Information Library of Alexandria (LILA) for species not included in the NACTI dataset. Additionally, we also included images provided for an important exotic species (nilgai; Kutugata et al. 2021) and for several arctic species (arctic fox, arctic wolves, polar bear, and caribou; R. Brook unpublished data). Our final training dataset contained images for 168 North American species. A complete list of images by species and how bounding boxes were generated is provided in the supplemental information.

### Model training

Prior to training models, we split the dataset into training and validation sets, with 70% used for training and 30% used for validation using the Python library scikit-learn v.0.23.2 (Pedregosa et al. 2011). Images were resized to a width and height of 408 and 307 pixels, respectively prior to training. We trained models using the Faster R-CNN architecture with a ResNet-50 feature pyramid network backbone (Ren et al. 2016) using the Pytorch v.1.9.1 library (Paszke et al. 2019) in Python v.3.8.5 (Python Software Foundation 2020). We obtained the architecture with weights that were pre-trained on the COCO (Common Objects in Context) dataset (Lin et al. 2015) from torchvision v.0.9.0 (Marcel and Rodriguez 2010). We used stochastic gradient descent with momentum for the optimizer, with a momentum value of 0.9 and a weight decay rate of 0.0005. The learning rate changed throughout training by using a plateau-based learning rate scheduler that began with an initial learning rate of 0.005 and reduced the learning rate by a factor of 0.5 once the validation loss stopped improving. We set initial anchor box sizes at 16, 32, 64, 128, and 256, with aspect ratios of 0.5, 1, and 2.

The model was trained with a batch size of 16 images. In the first epoch, the pre-trained weights from the COCO dataset were used and the best loss value was set to infinity (ensuring that subsequent epochs would have improved loss by comparison). After every epoch, the model was deployed on the validation dataset and the validation loss was calculated. If the validation loss was less than the best loss value from any epoch, the weights were saved and loaded in the next epoch. If the validation loss was worse than the best loss value, the weights were discarded and the best weights were loaded in the next epoch.

We trained models representing three levels of taxonomic classification. One is a taxonomic class level model that recognizes mammals (human and non-human) and birds. The second model classifies animals to taxonomic family for 31 families within mammals, birds, and reptiles. The third model is specific to North American fauna and recognizes 75 species of mammals, birds, and reptiles. All models include a category for vehicles and empty images. For the species and family models, we only included categories if there were at least 300 examples available for training and validation. See Tables 1-3 for details of what is included in each model.

**Table 1:** Performance metrics for the general model on validation data.

| Class | Precision | Recall | F1 | Average<br>Precision |
| --- | --- | --- | --- | --- |
| Aves | 0.906 | 0.995 | 0.948 | 0.964 |
| Human | 0.971 | 0.998 | 0.984 | 0.987 |
| Mammal | 0.927 | 0.992 | 0.958 | 0.948 |
| Vehicle | 0.971 | 0.993 | 0.982 | 0.959 |
| Mean | 0.944 | 0.994 | 0.968 | 0.965 |

**Table 2:**
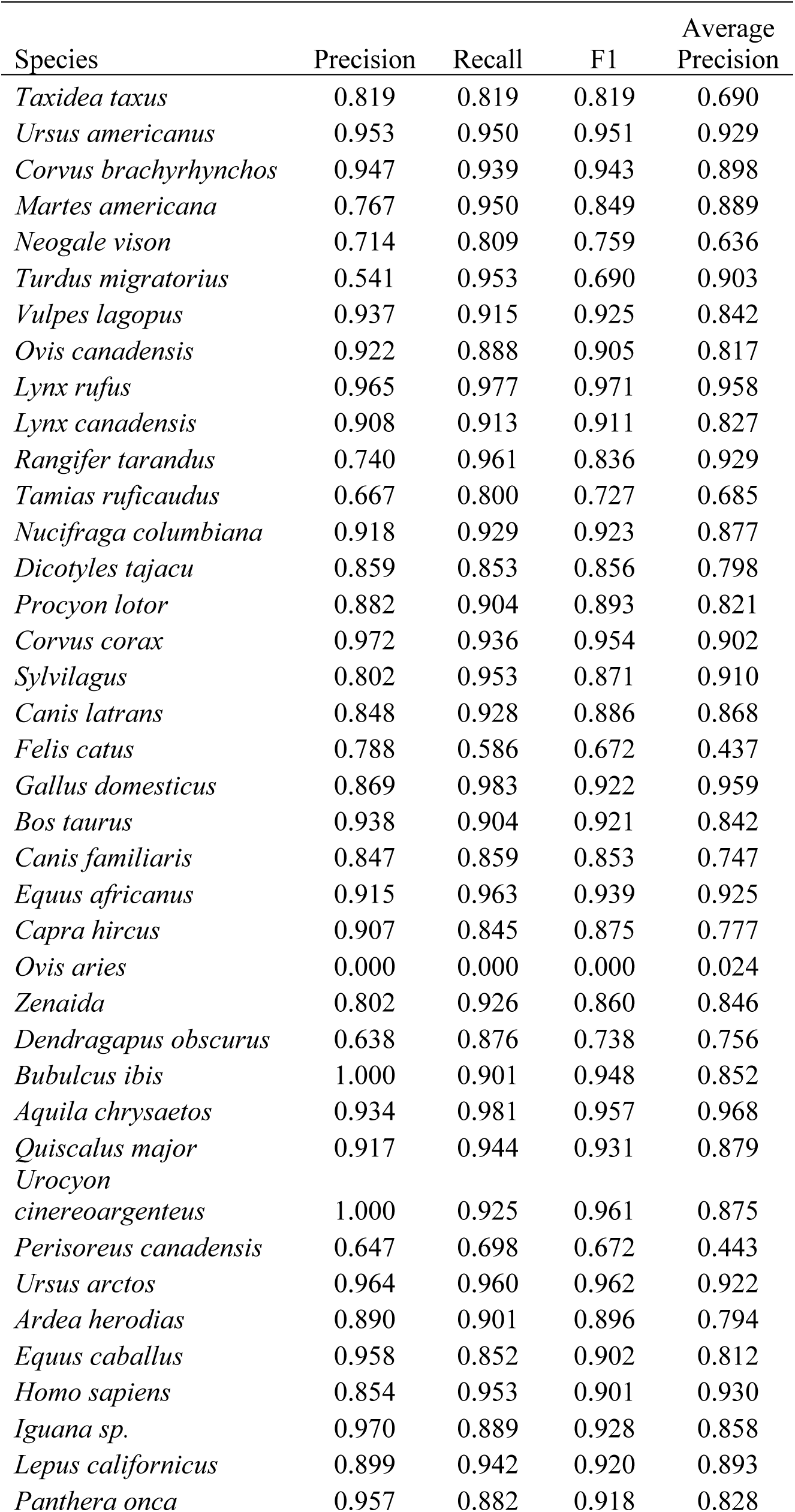
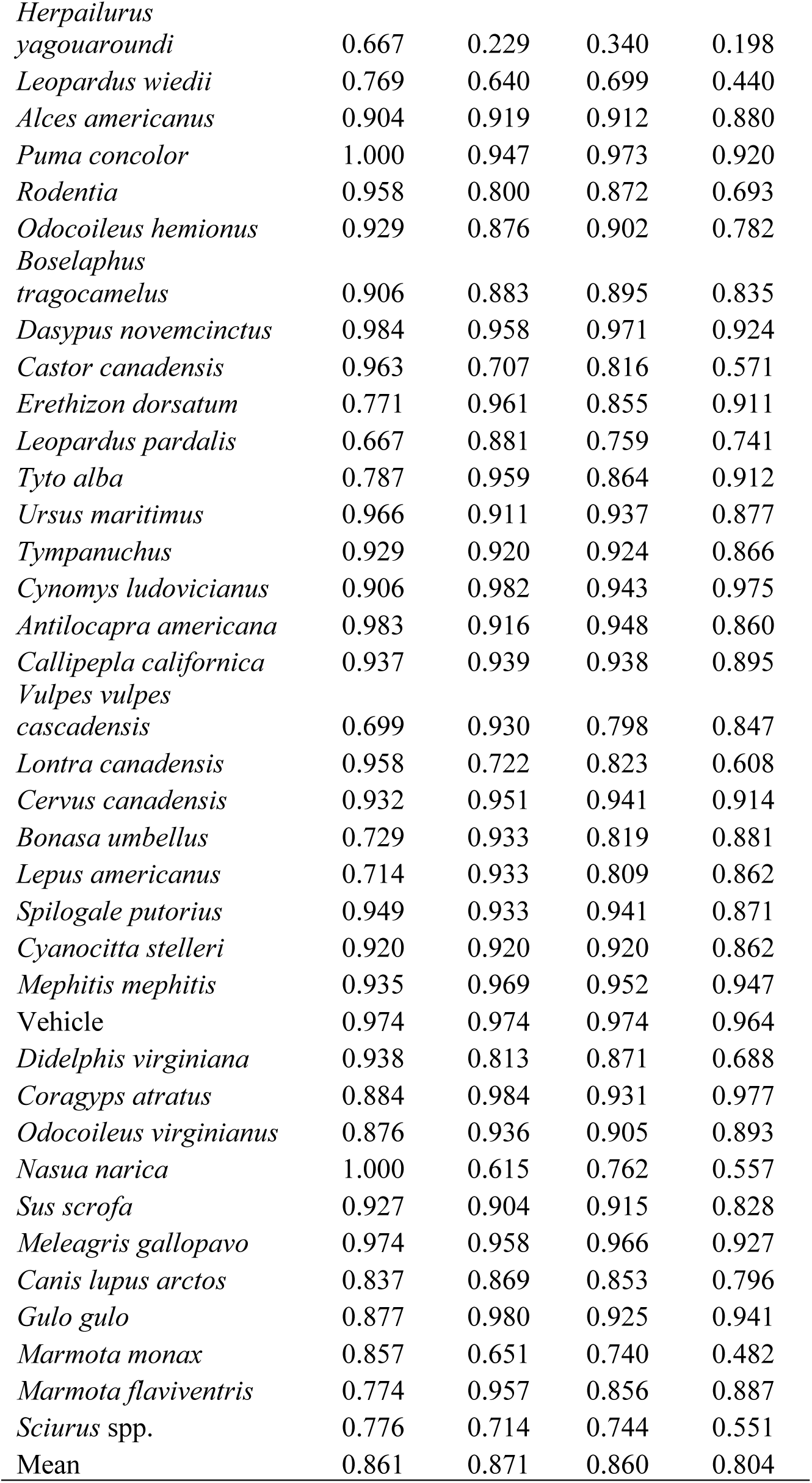
Performance metrics for the species model on validation data.

**Table 3:**
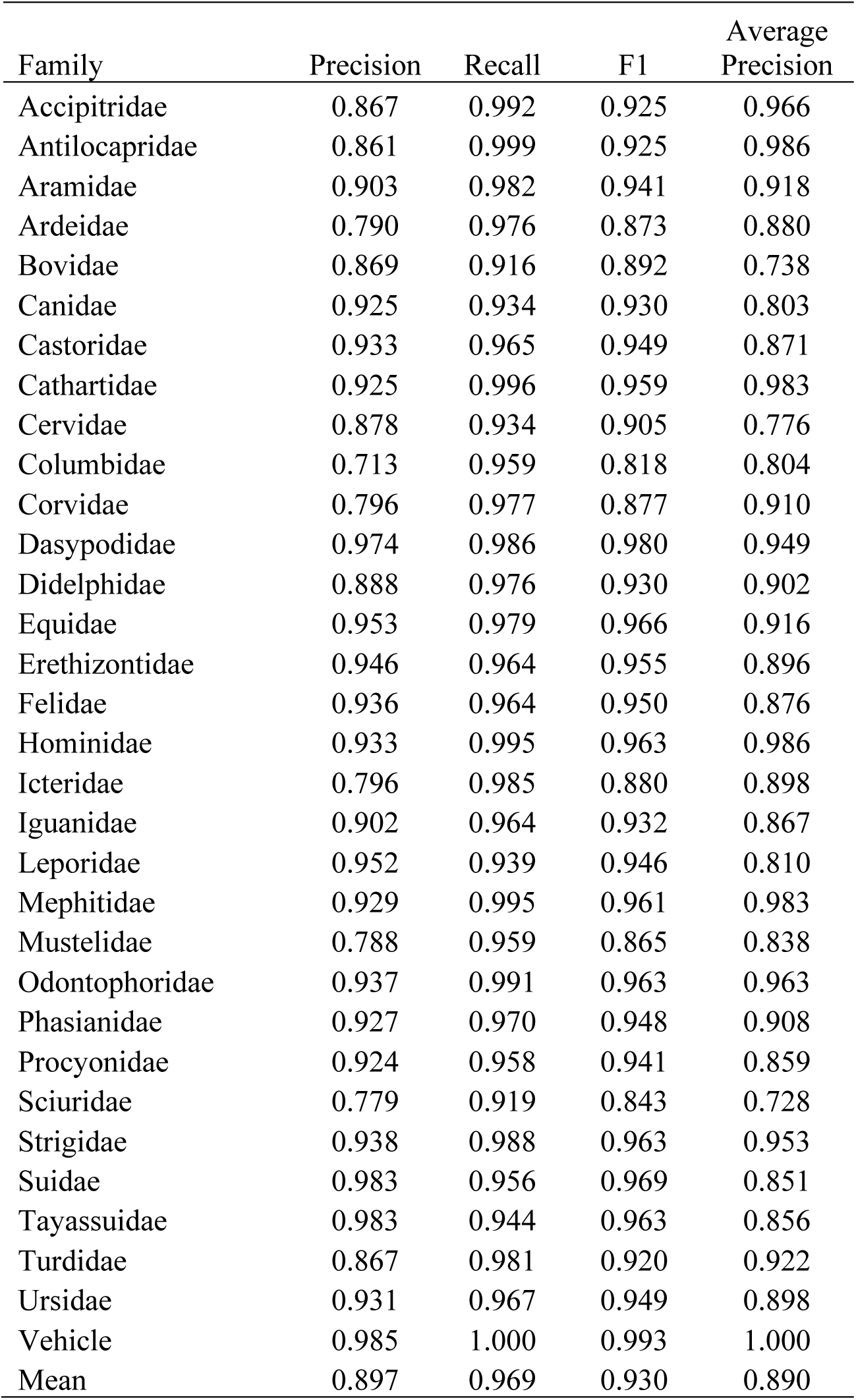
Performance metrics for the family model on validation data.

| Family | Precision | Recall | F1 | Average Precision |
| --- | --- | --- | --- | --- |
| Accipitridae | 0.867 | 0.992 | 0.925 | 0.966 |
| Antilocapridae | 0.861 | 0.999 | 0.925 | 0.986 |
| Aramidae | 0.903 | 0.982 | 0.941 | 0.918 |
| Ardeidae | 0.790 | 0.976 | 0.873 | 0.880 |
| Bovidae | 0.869 | 0.916 | 0.892 | 0.738 |
| Canidae | 0.925 | 0.934 | 0.930 | 0.803 |
| Castoridae | 0.933 | 0.965 | 0.949 | 0.871 |
| Cathartidae | 0.925 | 0.996 | 0.959 | 0.983 |
| Cervidae | 0.878 | 0.934 | 0.905 | 0.776 |
| Columbidae | 0.713 | 0.959 | 0.818 | 0.804 |
| Corvidae | 0.796 | 0.977 | 0.877 | 0.910 |
| Dasypodidae | 0.974 | 0.986 | 0.980 | 0.949 |
| Didelphidae | 0.888 | 0.976 | 0.930 | 0.902 |
| Equidae | 0.953 | 0.979 | 0.966 | 0.916 |
| Erethizontidae | 0.946 | 0.964 | 0.955 | 0.896 |
| Felidae | 0.936 | 0.964 | 0.950 | 0.876 |
| Hominidae | 0.933 | 0.995 | 0.963 | 0.986 |
| Icteridae | 0.796 | 0.985 | 0.880 | 0.898 |
| Iguanidae | 0.902 | 0.964 | 0.932 | 0.867 |
| Leporidae | 0.952 | 0.939 | 0.946 | 0.810 |
| Mephitidae | 0.929 | 0.995 | 0.961 | 0.983 |
| Mustelidae | 0.788 | 0.959 | 0.865 | 0.838 |
| Odontophoridae | 0.937 | 0.991 | 0.963 | 0.963 |
| Phasianidae | 0.927 | 0.970 | 0.948 | 0.908 |
| Procyonidae | 0.924 | 0.958 | 0.941 | 0.859 |
| Sciuridae | 0.779 | 0.919 | 0.843 | 0.728 |
| Strigidae | 0.938 | 0.988 | 0.963 | 0.953 |
| Suidae | 0.983 | 0.956 | 0.969 | 0.851 |
| Tayassuidae | 0.983 | 0.944 | 0.963 | 0.856 |
| Turdidae | 0.867 | 0.981 | 0.920 | 0.922 |
| Ursidae | 0.931 | 0.967 | 0.949 | 0.898 |
| Vehicle | 0.985 | 1.000 | 0.993 | 1.000 |
| Mean | 0.897 | 0.969 | 0.930 | 0.890 |

### Model evaluation

Once the model was trained, we evaluated its performance on the validation dataset. We calculated mean average precision (mAP) following standard methods for evaluating object detection algorithms (Girshick et al. 2016). For each category, a true positive (TP) is defined as a match between a bounding box in the ground truth annotation and the prediction. Specifically, the Intersection over Union (IoU) must be greater than or equal to 0.5, where:

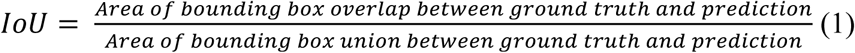

A false positive (FP) occurs when a predicted bounding box occurs in an image either without a ground truth bounding box for that species or the predicted bounding box has an IoU with the ground truth bounding box is less than 0.5. A false negative (FN) occurs when there is a ground truth bounding box that does not have a matching predicted bounding box with an IoU ≥ 0.5. Then, we calculated recall and precision for each category, where:

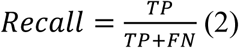

and

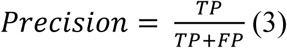

For each category we calculated the average precision, which is a measure of the area under the curve from a plot with recall on the x-axis and precision on the y-axis. Specifically, for a series of confidence scores (10 values equally spaced between zero and one), we calculated recall and precision for each confidence score, plotted recall by precision and calculated the area under the curve. Mean average precision is the mean of the average precision across all categories.

While mAP is a standard metric for object detection models, many ecologists are less concerned about the predicted location of the bounding box and more concerned about the predicted contents of the image and the ability to accurately identify empty images (those without animals or vehicles). Therefore, we also calculated detection-level precision, recall, and F_1_ score, where

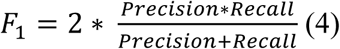

for each category at the confidence score threshold that we recommend using in CameraTrapDetectoR (0.5). Detection-level metrics were calculated differently from how they are calculated for image classification models, as there can be multiple classifications in each image (Zhang and Zhou 2014). For each category in each image, TPs are determined if the number of objects predicted matches the number observed. If more objects are predicted than observed, these are FPs, and if fewer objects are predicted than observed, these are FNs. We used these rates to calculate the metrics for each category following equations 2-4.

### R package development

CameraTrapDetectoR was developed in R v.4.1.2 (R Core Team 2021), and works with several types of image files including JPG, TIF, PNG, or PDF. The package magick v.2.7.3 (Ooms 2021) is used for reading and manipulating images. Our package uses the R package torch v.0.6.0 (Falbel et al. 2021) to deploy trained models and torchvision v.0.4.0 (Falbel 2021) for converting images to tensors, which are the unit upon which torch deploys models. Roxygen2 v.7.1.2 (Wickham et al. 2021) was used to generate manual files. We developed Shiny Applications using the R package Shiny v.1.7.1 (Chang et al. 2021) and shinyFiles v.0.9.1 (Pedersen et al. 2021).

## Results

### Model performance on validation dataset

All models performed well on the validation datasets (Tables 1-3; Figures 1-2). Mean average precision was the highest for the taxonomic class model (0.96) and lowest for the species model (0.80), with the taxonomic family model having an intermediate mAP value (0.89). Category-level recall, precision and F_1_ scores followed the same trend of the class model performing best and species model performing worst (Tables 1-3). The distributions of detection confidence scores tended to be much higher for accurately identified animals versus misclassifications (Figure 3 and Figures S1-3). For categories with the highest precision the distributions had the largest difference.

**Figure 1.**
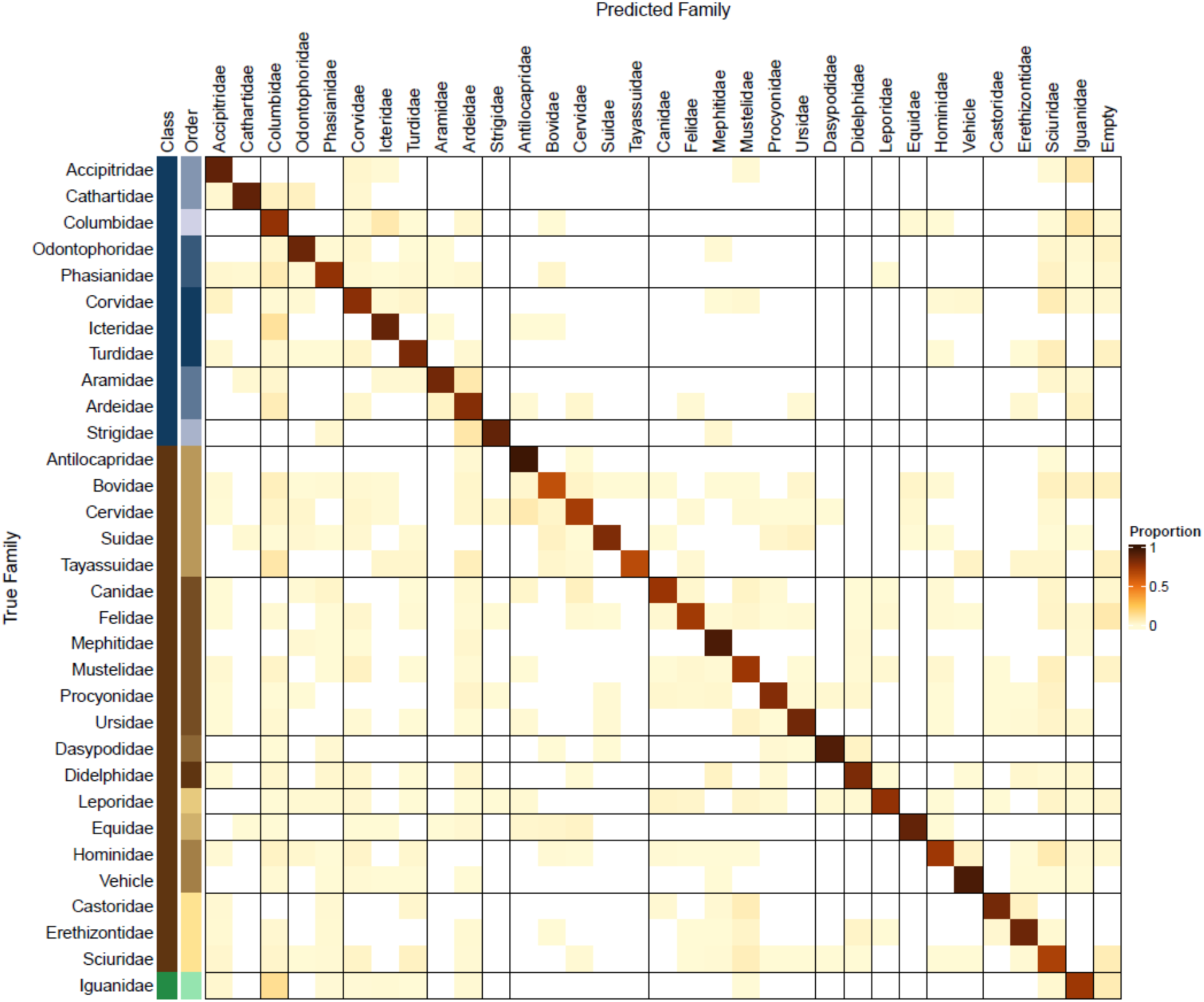
Confusion matrix for 31 avian (blue), mammal (brown) and reptile (green) taxonomic families. Cell color represents the proportion of images correctly classified (diagonal cells) and images misclassified (off diagonal cells). Vertical and horizonal lines indicate taxonomic order. Misclassification was low for all taxonomic families – indicated by the pale yellow off-diagonal cells and dark brown diagonal cells.

**Figure 2.**
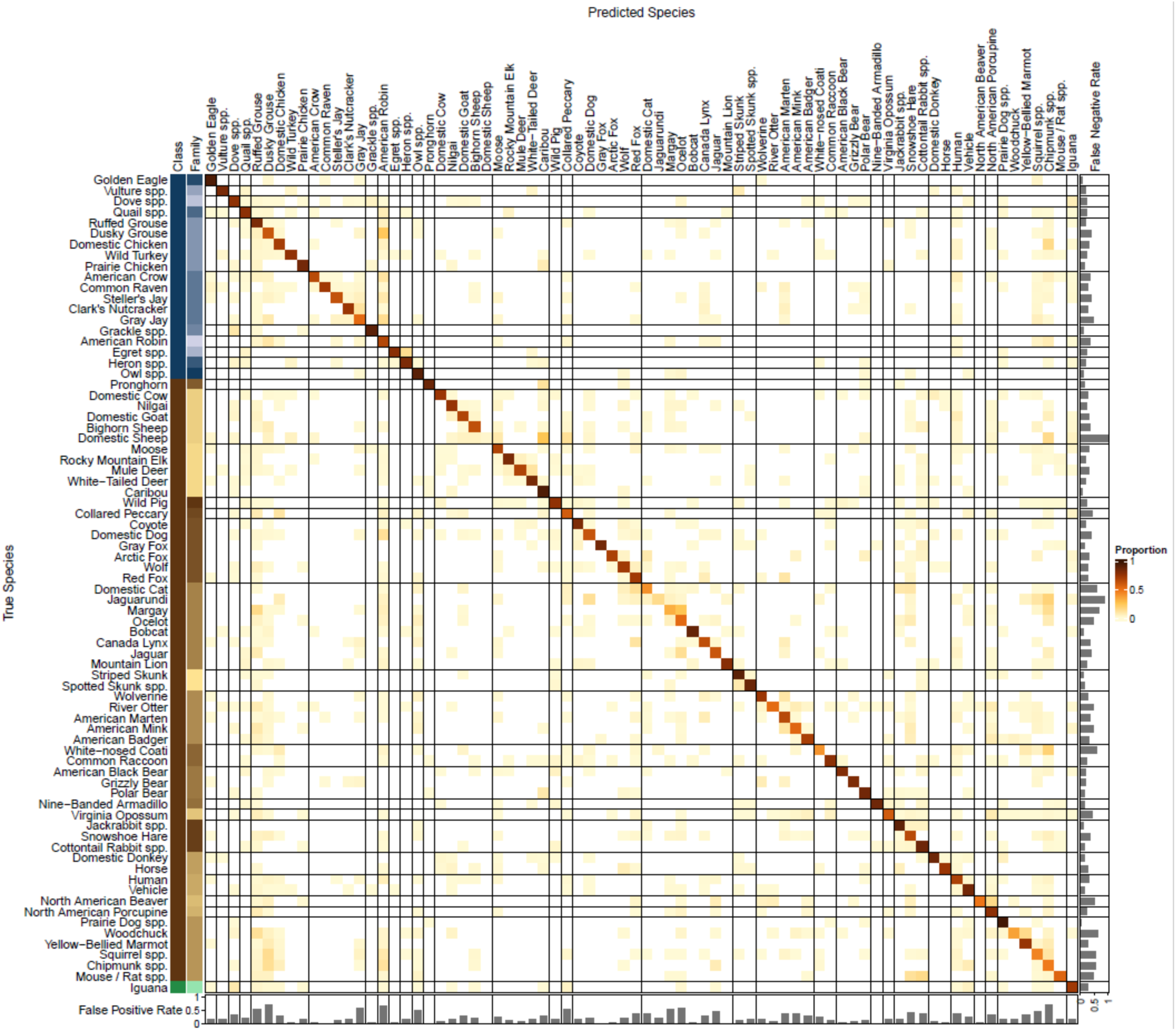
Confusion matrix for 75 avian (blue), mammal (brown) and reptile (green) species. Cell color represents the proportion of images correctly classified (diagonal cells) and images misclassified (off diagonal cell). Vertical and horizonal lines indicate taxonomic families. Misclassification was generally low for all species – indicated by the pale yellow off-diagonal cells.

**Figure 3.**
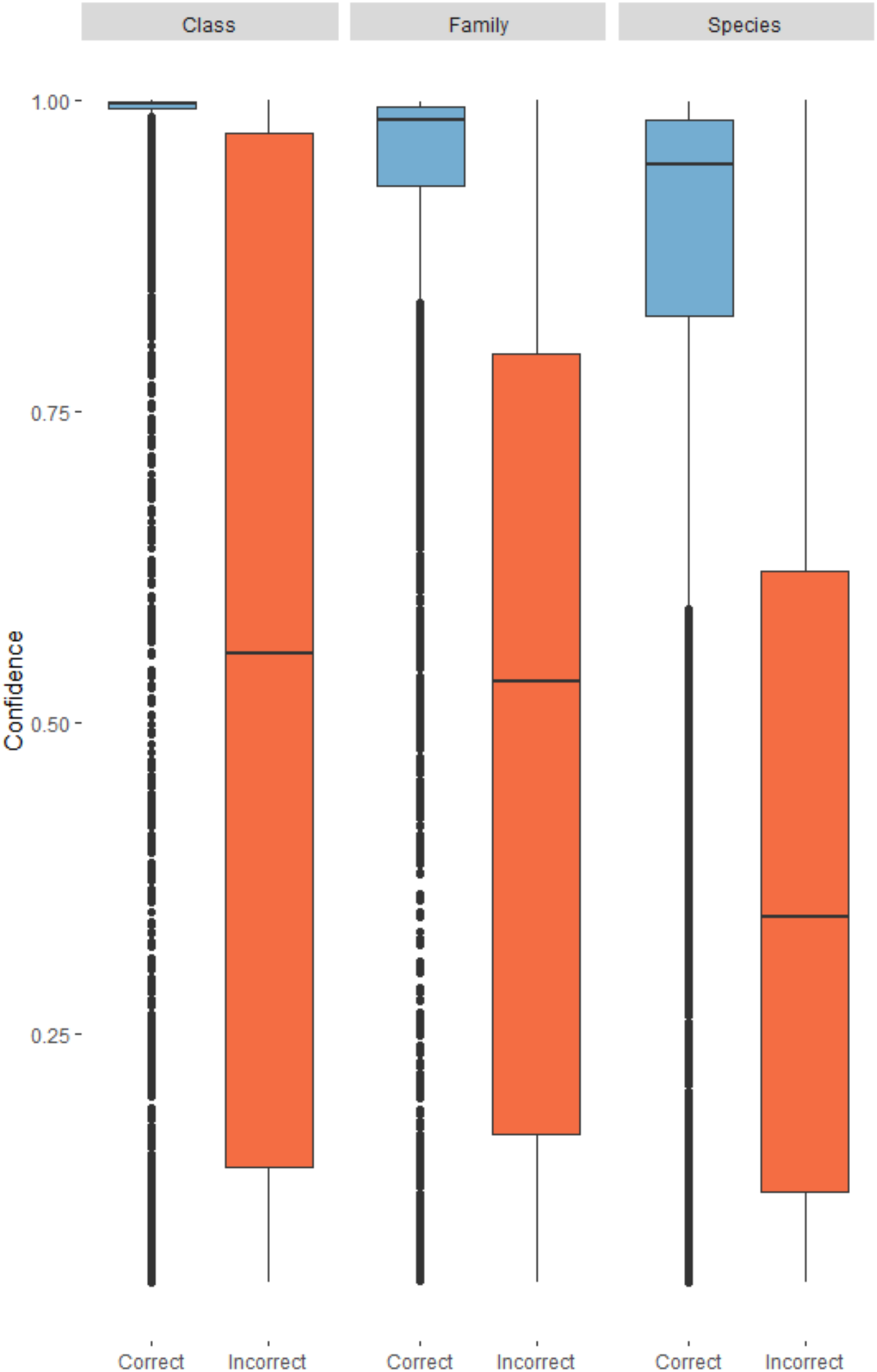
Distribution of confidence scores for correctly identified (blue) and incorrectly (orange) identified categories in all three models.

## Discussion

Our models are effective at identifying and counting species of interest and at distinguishing empty images from those with animals. There are several software options available for automatically processing camera trap images using deep learning computer vision algorithms (Vélez et al. 2022). Our R package is most similar in functionality to MegaDetector (Beery et al. 2019), a powerful and effective algorithm that detects, classifies, and counts humans, non-human animals, and vehicles in camera trap images. MegaDetector offers users the ability run locally on their computers by running Python scripts, or users can send images to the development team to have them analyzed. The output is a JSON file depicting the coordinates and predictions of bounding boxes for objects in each image.

CameraTrapDetectoR runs directly in R and does not require a Python installation. From comments about previous R packages built by some of our team (Tabak et al. 2019, 2020b), many biologists prefer avoiding the Python language for various reasons. Our package will also, optionally, create plots of each image in a dataset with the predicted bounding box and category label (e.g., Figure 4). This option allows users to evaluate model performance in real time as the model is deploying. CameraTrapDetectoR is developed specifically for Windows operating systems, which avoids some of the other issues that users have had with previous R packaged developed by our team. Support for Macintosh and Linux operating systems will be available in the near future.

**Figure 4.**
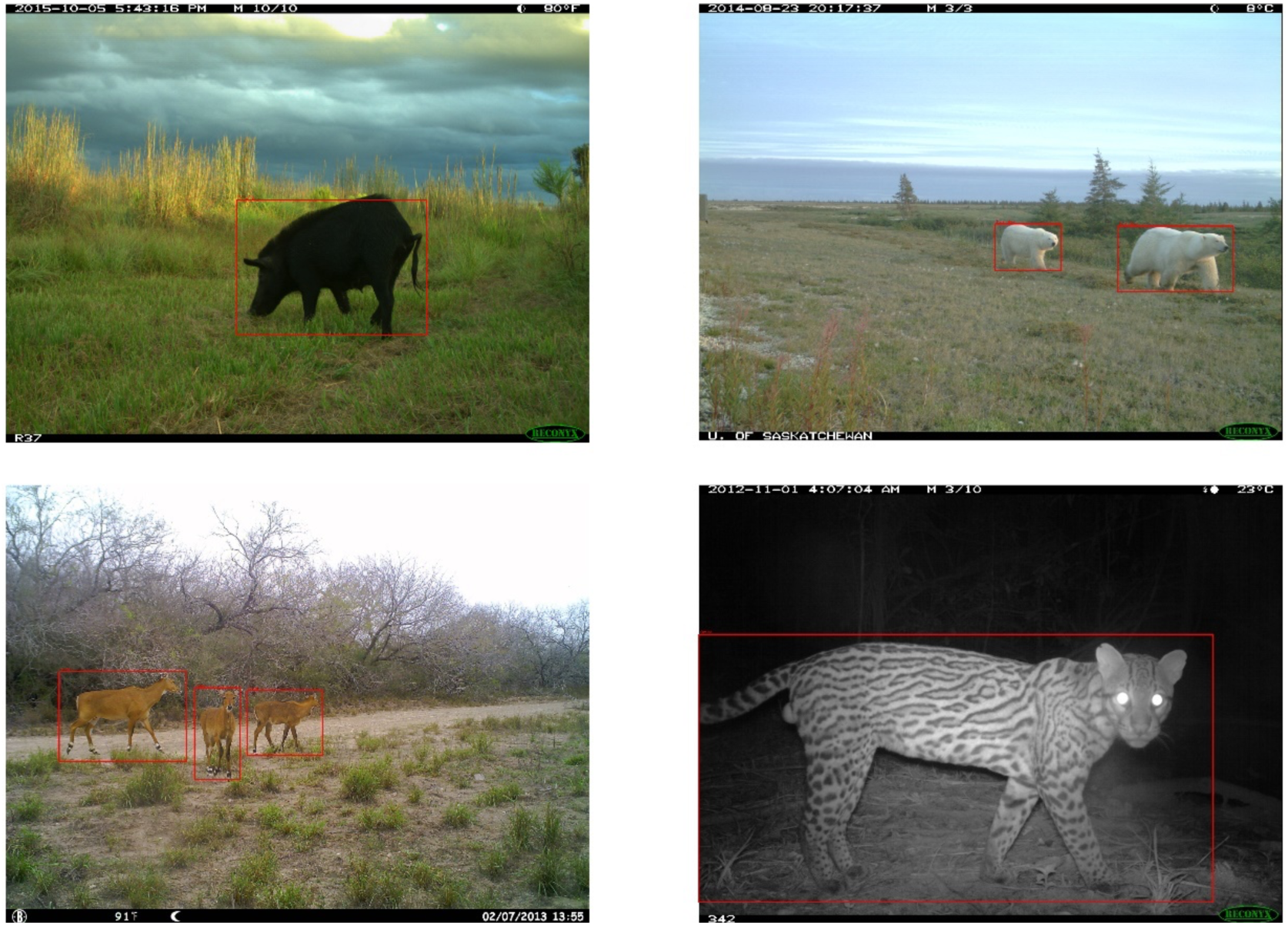
Example output from CameraTrapDetectoR when classifying images using the species model. When deploying models, users can optionally have predicted bounding boxes plotted with the original image. When multiple animals occur in the image, bounding boxes are generated for each animal identified providing a count of the total number of animals by species for each image.

CameraTrapDetectoR offers four main advantages when compared to the currently available options: 1) our package uses a taxonomic approach including models for three taxonomic levels – class, family, and species. This is especially useful when the species of interest is not included in the species level model. The taxonomic class model recognizes mammals (human and non-human) and birds. This model is effective at removing empty images from a wide variety of datasets and is similar to the model provided in MegaDetector. We also include a model that recognizes 75 common North American domestic and wild species, including all bear, wildcat and canid species, and a model that classifies these species to family, which can be useful for identifying species not included in the species level model. We found that our model was effective at finding these species in out-of-distribution datasets.

2)Frequently camera trap image classification is done as part of a larger workflow. CameraTrapDetectoR can be integrated into automated or semi-automated work flows to speed image processing. This allows for images to be processed in near real time allowing for improved monitoring of populations.

3)Frequently there are privacy or intellectual property concerns that prevent camera trap images from being shared. CameraTrapDetectoR runs locally on laptop computers and requires very minimal programming. After installing the R package, a user only needs to run one line of code to launch a Shiny Application that will facilitate model deployment using a point-and-click interface.

4)After deploying the model, predictions are written to a CSV file where the number of animals in each category are predicted for each image file; this information is also saved as an R data frame. We think that the tabular output format are more intuitive to interpret than a JSON file, and can be directly incorporated into analysis pipelines. As several R packages can be used for estimating occupancy and abundance of animals based on detections from camera trap data (Fiske and Chandler 2011, Oksanen 2020, Tabak et al. 2020a), one could pair these packages with CameraTrapDetectoR to automatically generate estimates of animal status from camera trap images.

Some species and families were more difficult to accurately identify - specifically smaller wildcat, domestic cats, smaller avian species, and rodents (Figure 2). For some of these species it is likely due to smaller training datasets. Species such as ocelot, margay, jaguarundi, and domestic cat may require very large training datasets to accurately differentiate due to similar pillage and size. When an image was misclassified, it was most commonly misclassified as smaller species of Galliformes, Sciuridae, *Lepus*, or *Sylvilagus*. This may be due to a combination of number of images available for training as well as the size of the animals in the image is generally smaller. Our package can help users with categories for which our model performed poorly by providing the option to specify the confidence score threshold. If users are interested in ensuring detection of these species for which our models performed poorly, a potential approach is to set the confidence score threshold low (e.g. score_threshold=0.1). This will result in fewer false negatives. If users are concerned about obtaining too many false positives, they should set the confidence score threshold to a higher value (e.g., score_threshold=0.8). Furthermore, by specifying write_bbox_csv=TRUE, the package will produce a csv file with all predicted bounding boxes, regardless of confidence score (and confidence score is included in this csv).

In CameraTrapDetectoR, we provide an easy and effective means for detecting, identifying, and counting animals in camera trap images. The ease of operating this object detection system in R can facilitate rapid deployment and results in near real-time. For example, if researchers deploy camera traps that relay images by satellite to a computer (Whytock et al. 2021), the computer can automatically deploy the models in our package in R and provide results. This can be especially valuable for monitoring invasive species and species of conservation concern.

## References

Beery, S., D. Morris, and S. Yang. 2019. Efficient Pipeline for Camera Trap Image Review. 1907.06772 [cs].

Beery, S., G. Wu, V. Rathod, R. Votel, and J. Huang. 2020. Context R-CNN: Long Term Temporal Context for Per-Camera Object Detection. Pages 13075–13085.

Chalmers, C., P. Fergus, S. Wich, and A. C. Montanez. 2019. Conservation AI: Live Stream Analysis for the Detection of Endangered Species Using Convolutional Neural Networks and Drone Technology. 1910.07360 [cs].

Chang, W., J. Cheng, J. J. Allaire, C. Sievert, B. Schloerke, Y. Xie, J. Allen, J. McPherson, A. Dipert, and B. Borges. 2021. shiny: Web Application Framework for R.

Falbel, D. 2021. torchvision: Models, Datasets and Transformations for Images.

Falbel, D., J. Luraschi, D. Selivanov, A. Damiani, C. Regouby, K. Joachimiak, and RStudio. 2021. torch: Tensors and Neural Networks with “GPU” Acceleration.

Falzon, G., C. Lawson, K.-W. Cheung, K. Vernes, G. A. Ballard, P. J. S. Fleming, A. S. Glen, H. Milne, A. Mather-Zardain, and P. D. Meek. 2020. ClassifyMe: A Field-Scouting Software for the Identification of Wildlife in Camera Trap Images. Animals 10:58.

Fiske, I., and R. Chandler. 2011. unmarked : An R Package for Fitting Hierarchical Models of Wildlife Occurrence and Abundance. Journal of Statistical Software 43.

Girshick, R., J. Donahue, T. Darrell, and J. Malik. 2016. Region-Based Convolutional Networks for Accurate Object Detection and Segmentation. IEEE Transactions on Pattern Analysis and Machine Intelligence 38:142–158.

He, K., X. Zhang, S. Ren, and J. Sun. 2016. Deep Residual Learning for Image Recognition. Pages 770–778 2016 IEEE Conference on Computer Vision and Pattern Recognition (CVPR). IEEE, Las Vegas, NV, USA.

Jumeau, J., L. Petrod, and Y. Handrich. 2017. A comparison of camera trap and permanent recording video camera efficiency in wildlife underpasses. Ecology and Evolution 7:7399–7407.

Kutugata, M., J. Baumgardt, J. A. Goolsby, and A. E. Racelis. 2021. Automatic Camera-Trap Classification Using Wildlife-Specific Deep Learning in Nilgai Management. Journal of Fish and Wildlife Management 12:412–421.

Lin, T.-Y., M. Maire, S. Belongie, L. Bourdev, R. Girshick, J. Hays, P. Perona, D. Ramanan, C. L. Zitnick, and P. Dollár. 2015. Microsoft COCO: Common Objects in Context. 1405.0312 [cs].

Marcel, S., and Y. Rodriguez. 2010. Torchvision the machine-vision package of torch. Pages 1485–1488 Proceedings of the 18th ACM international conference on Multimedia. Association for Computing Machinery, New York, NY, USA.

Niedballa, J., R. Sollmann, A. Courtiol, and A. Wilting. 2016. camtrapR: an R package for efficient camera trap data management. Methods in Ecology and Evolution 7:1457–1462.

Norouzzadeh, M. S., D. Morris, S. Beery, N. Joshi, N. Jojic, and J. Clune. 2021. A deep active learning system for species identification and counting in camera trap images. Methods in Ecology and Evolution 12:150–161.

Norouzzadeh, M. S., A. Nguyen, M. Kosmala, A. Swanson, M. S. Palmer, C. Packer, and J. Clune. 2018. Automatically identifying, counting, and describing wild animals in camera-trap images with deep learning. Proceedings of the National Academy of Sciences 115:E5716–E5725.

O’Connor, K. M., L. R. Nathan, M. R. Liberati, M. W. Tingley, J. C. Vokoun, and T. A. G. Rittenhouse. 2017. Camera trap arrays improve detection probability of wildlife: Investigating study design considerations using an empirical dataset. PLOS ONE 12:e0175684.

Oksanen, J. 2020. Vegan: ecological diversity:12.

Ooms, J. 2021. magick: Advanced Graphics and Image-Processing in R.

Paszke, A., S. Gross, F. Massa, A. Lerer, J. Bradbury, G. Chanan, T. Killeen, Z. Lin, N. Gimelshein, L. Antiga, A. Desmaison, A. Kopf, E. Yang, Z. DeVito, M. Raison, A. Tejani, S. Chilamkurthy, B. Steiner, L. Fang, J. Bai, and S. Chintala. 2019. PyTorch: An Imperative Style, High-Performance Deep Learning Library. Pages 8026–8037 in H. Wallach, H. Larochelle, A. Beygelzimer, F. Alché-Buc, E. Fox, and R. Garnett, editors. Advances in Neural Information Processing Systems 32. Curran Associates, Inc.

Pedersen, T. L., V. Nijs, T. Schaffner, and E. Nantz. 2021. shinyFiles: A Server-Side File System Viewer for Shiny.

Pedregosa, F., G. Varoquaux, A. Gramfort, V. Michel, B. Thirion, O. Grisel, M. Blondel, P. Prettenhofer, R. Weiss, V. Dubourg, J. Vanderplas, A. Passos, D. Cournapeau, M. Brucher, M. Perrot, and É. Duchesnay. 2011. Scikit-learn: machine learning in python. Journal of Machine Learning Research 12:2825–2830.

Price Tack, J. L., B. S. West, C. P. McGowan, S. S. Ditchkoff, S. J. Reeves, A. C. Keever, and J. B. Grand. 2016. AnimalFinder: A semi-automated system for animal detection in time-lapse camera trap images. Ecological Informatics 36:145–151.

Python Software Foundation. 2020. Python. Beaverton, OR.

R Core Team. 2021. R: A language and Environment for Statistical Computing. Vienna, Austria.

Ren, S., K. He, R. Girshick, and J. Sun. 2016. Faster R-CNN: Towards Real-Time Object Detection with Region Proposal Networks. 1506.01497 [cs].

Schneider, S., S. Greenberg, G. W. Taylor, and S. C. Kremer. 2020. Three critical factors affecting automated image species recognition performance for camera traps. Ecology and Evolution 10:3503–3517.

Sollmann, R. 2018. A gentle introduction to camera-trap data analysis. African Journal of Ecology 56:740–749.

Swanson, A., M. Kosmala, C. Lintott, R. Simpson, A. Smith, and C. Packer. 2015. Snapshot Serengeti, high-frequency annotated camera trap images of 40 mammalian species in an African savanna. Scientific Data 2:150026.

Tabak, M. A., J. S. Lewis, P. E. Schlichting, N. P. Snow, K. C. VerCauteren, and R. S. Miller. 2020a. rapidPop: Rapid population assessments of wildlife using camera trap data in R Shiny Applications. Page 2020.03.30.017103.

Tabak, M. A., M. S. Norouzzadeh, D. W. Wolfson, E. J. Newton, R. K. Boughton, J. S. Ivan, E. A. Odell, E. S. Newkirk, R. Y. Conrey, J. Stenglein, F. Iannarilli, J. Erb, R. K. Brook, A. J. Davis, J. Lewis, D. P. Walsh, J. C. Beasley, K. C. VerCauteren, J. Clune, and R.S. Miller. 2020b. Improving the accessibility and transferability of machine learning algorithms for identification of animals in camera trap images: MLWIC2. Ecology and Evolution 10:10374–10383.

Tabak, M. A., M. S. Norouzzadeh, D. W. Wolfson, S. J. Sweeney, K. C. Vercauteren, N. P. Snow, J. M. Halseth, P. A. D. Salvo, J. S. Lewis, M. D. White, B. Teton, J. C. Beasley, P. E. Schlichting, R. K. Boughton, B. Wight, E. S. Newkirk, J. S. Ivan, E. A. Odell, R. K. Brook, P. M. Lukacs, A. K. Moeller, E. G. Mandeville, J. Clune, and R. S. Miller. 2019. Machine learning to classify animal species in camera trap images: Applications in ecology. Methods in Ecology and Evolution 10:585–590.

Vélez, J., P. J. Castiblanco-Camacho, M. A. Tabak, C. Chalmers, P. Fergus, and J. Fieberg. 2022. Choosing an Appropriate Platform and Workflow for Processing Camera Trap Data using Artificial Intelligence. 2202.02283 [cs].

Whytock, R. C., T. Suijten, T. van Deursen, J. Świeżewski, H. Mermiaghe, N. Madamba, N. Moukoumou, J. A. Zwerts, A. F. K. Pambo, L. Bahaa-el-din, S. Brittain, A. W. Cardoso, P. Henschel, D. Lehmann, B. R. Momboua, L. Makaga, C. Orbell, L. J. T. White, D. M. Iponga, and K. A. Abernethy. 2021. Real-time alerts from AI-enabled camera traps using the Iridium satellite network: a case-study in Gabon, Central Africa. Page 2021.11.10.468078.

Wickham, H., P. Danenberg, G. Csárdi, M. Eugster, and RStudio. 2021. roxygen2: In-Line Documentation for R.

Wildlife Insights. 2021. Wildlife insights artificial intelligence models.

Willi, M., R. T. Pitman, A. W. Cardoso, C. Locke, A. Swanson, A. Boyer, M. Veldthuis, and L. Fortson. 2019. Identifying animal species in camera trap images using deep learning and citizen science. Methods in Ecology and Evolution 10:80–91.

Yousif, H., R. Kays, and H. Zhihai. 2019. Dynamic programming selection of object proposals for sequence-level animal species classification in the wild. IEEE Transactions on Circuits and Systems for Video Technology.

Zhang, M.-L., and Z.-H. Zhou. 2014. A Review On Multi-Label Learning Algorithms. Knowledge and Data Engineering, IEEE Transactions on 26:1819–1837.

Zhang, Z., Z. He, G. Cao, and W. Cao. 2016. Animal detection from highly cluttered natural scenes using spatiotemporal object region proposals and patch verification. IEEE Transactions on Multimedia 18:2079–2092.

Zou, Z., Z. Shi, Y. Guo, and J. Ye. 2019. Object Detection in 20 Years: A Survey. 1905.05055 [cs].

